# The gift of novelty: repeat-robust *k*-mer-based estimators of mutation rates

**DOI:** 10.64898/2026.04.01.715966

**Authors:** Haonan Wu, Paul Medvedev

## Abstract

Estimating mutation rates between evolutionarily related sequences is a central problem in molecular evolution. Due to the rapid expansion of datasets, modern methods avoid costly alignment and instead focus on comparing sketches of sets of constituent *k*-mers. While these methods perform well on many sequences, they are not robust to highly repetitive sequences such as centromeres. In this paper, we present three new estimators that are robust to the presence of repeats. The estimators are applicable in different settings, based on whether they need count information from zero, one, or both of the sequences. We evaluate our estimators empirically using highly repetitive alpha satellite sequences. Our estimators each perform best in their class and our strongest estimator outperforms all other tested estimators. Our software is open-source and freely available on https://github.com/medvedevgroup/Accurate_repeat-aware_kmer_based_estimator.

## 1. Introduction

Estimating mutation rates between evolutionarily related sequences has long been a central problem in molecular evolution, originating well before the advent of large-scale genomics. Early quantitative methods concentrated on amino acid substitution rates, such as the PAM matrices introduced by Dayhoff (1969) and the BLOSUM matrices developed by Henikoff and Henikoff (1992). These methodologies, along with later profile-based hidden Markov models (Durbin, 2013) continue to serve as the benchmark when high-quality alignments can be obtained.

Nevertheless, the rapid growth of sequencing has made computationally intensive alignment-based pipelines increasingly infeasible in the modern era. As a result, alignment-free methods that characterize sequences using low-cost summary statistics have become essential (Song et al., 2014; Zielezinski et al., 2017; Rathore and Kashyap, 2026). Most of these techniques are based on sketches of *k*-mer spectra. Widely used tools such as Mash (Ondov et al., 2016) and Skmer (Sarmashghi et al., 2019), along with more recent sketch-adjusted approaches including Sylph (Shaw and Yu, 2024) and FracMinHash-based methods (Hera et al., 2024, 2023; Hera and Koslicki, 2025; Hera et al., 2025; Shaw and Yu, 2023; Wu et al., 2025), enable rapid construction of whole-genome phylogenies, efficient metagenomic screening, and the estimation of millions of pairwise point-mutation rates in minutes rather than days.

Nearly all alignment-free approaches are derived based on the assumption that most *k*-mers above a certain *k*-mer size (e.g. *k* ≥ 19) occur only once in a sequence. However, recent advances in sequencing technology are leading to more abundantly available highly repetitive sequences. For example, the recent telomere-to-telomere human assembly contains fully assembled chromosome centromeres, which are alpha satellite DNA composed of 171-bp monomers that are further arranged into higher-order repeats (Logsdon et al., 2024). Unfortunately, most estimators are not robust to repeat-rich sequences and methods to analyze the mutation rates between such sequences remain in their infancy.

We can categorize the space of *k*-mer-based estimators based on the type of information they use. In the absence of repeats, it suffices to consider the set of *k*-mers present in the sequences and ignore their occurrence counts. We call these type of estimators as Presence-Presence, because they rely on presence/absence information for both the source string *s* and the mutated string *t*. Such estimators are especially useful in the setting where occurrence counts are not readily available, such as raw sequencing data. In the presence of repeats, however, occurrence counts become an important signal. In the Presence-Count setting, an estimator is restricted to presence/absence information for *s* but is allowed to use occurrence counts of *t*. This can occur if for example *s* is unassembled sequencing data while *t* is an assembly. The Count-Count setting is the most powerful, allowing the estimator to use information about counts in both *s* and *t*, but is limited to applications such as when both *s* and *t* are assembled.

The widely known Mash estimator falls in the Presence-Presence category, but, as we show in this paper, is not repeat-robust. There are two repeat-robust *k*-mer-based estimators that we are aware of. The first is from our previous work (Wu et al., 2025), which, as we will describe in Sec. 7, falls roughly in between the Presence-Presence and the Presence-Count setting. The second is a weighted-intersection-based estimator, which falls in the Count-Count setting. This is a natural estimator that has been mentioned in Rhie et al. (2020).

In this paper, we present three new estimators (Table 1): 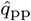, for the Presence-Presence setting, 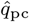, for the Presence-Count setting, and 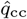, for the Count-Count setting. One of our main insights is that the number of newly created *k*-mers is more sensitive to the presence of repeats than the number of *k*-mers that remain shared (or, as reflected in the title, we treat *novel k*-mers as a *gift* to make use of).

**Table 1.** Our contributed estimators. Here, *s* in arbitrary string with *L* = |*s*| *− k* + 1; *t* is the result of applying a mutation process to *s* with substitution rate *r*; *d*_1_(*τ, s*) is the number of *k*-mers in the spectrum of *s* that are at a Hamming distance of one to *τ* . We use *q* as shorthand for 1 *−* (1 *− r*)^*k*^, e.g. an estimator 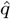 implicitly defines 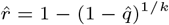 .

| Name | Knowledge of $k$-mers in $s$ | Knowledge of $k$-mers in $t$ | Formula |
| --- | --- | --- | --- |
| $\hat{q}_{pp}$ | Presence/absence | Presence/absence | $\frac{\vert sp(t) \setminus sp(s)\vert }{L}$ |
| $\hat{q}_{pc}$ | Presence/absence | Counts | $\sum_{\tau \in sp(t) \setminus sp(s)} \frac{occ(\tau, t)}{L}$ |
| $\hat{q}_{cc}$ | Counts | Counts | $\hat{q}_{pc} + (1 - \hat{r}_{pc})^{k-1} \cdot \frac{\hat{r}_{pc}}{3L} \cdot \sum_{\tau \in sp(s)} occ(\tau, s) \cdot d_1(\tau, s)$ |

We evaluate our estimators empirically using various types of sequences, including the alpha satellite centromeric region from a human chromosome. We evaluate estimator error across a wide range of mutation rates and *k*-mer values. Moreover, we demonstrate how our estimators can be combined with FracMinHash sketching without systematically effecting bias. Ultimately, each of our estimators performs best in their class, with 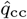 outperforming estimators in all classes. Finally, we show how our estimators can be used on real data by applying them to compute the ANI (Average Nucleotide Identity), which is widely used to measure genomic distance in taxonomic analysis.

## 2. Preliminaries

Let *s* be a string and let *k >* 0 be the *k*-mer size. We let |*s*| denote the number of nucleotides in *s*. We assume in this paper that |*s*| ≥ *k*. We use *L* to denote the number of *k*-mers in *s*, i.e. *L* = |*s*| − *k* + 1. Let *sp* (s) be the set of all distinct *k*-mers in *s*, also called the *spectrum* of *s*. Let *occ*(*τ, s*) denote the number of copies of *k*-mer *τ* in string *s*. Let *d*_*i*_(*τ, s*) denote the number of *k*-mers in *sp*(*s*) with Hamming distance *i* to *k*-mer *τ* . The Jaccard similarity between two sets *A* and *B* is 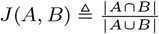 .

We will consider the simple substitution mutation model as in previous work (Blanca et al., 2022). Given a parameter 0 ≤ *r* ≤ 1 and a string *s*, the model generates an equal-length string where, independently, the character at each position is unchanged from *s* with probability 1 − *r* and changed to one of the three other nucleotides with probability *r/*3.

Our goal in this paper is to estimate the mutation rate *r* given observations about *s* and *t*. We assume that, at a minimum, we can observe *L, sp*(*s*), and *sp*(*t*). Depending on the category of the estimator, we may also observe *occ*(*τ, s*) and/or *occ*(*τ, t*), for all *τ* .

As a shorthand notation, we define *q* ≜ 1 − (1 − *r*)^*k*^; intuitively, *q* is the probability that a *k*-mer is mutated. In this work and others, estimators for *r* are usually derived by first obtaining an estimator 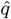 for *q* and then computing the estimator 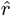 using the inverse formula 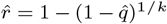. We take the same approach in this paper, where we will explicitly define an estimator for *q* and leave the definition of *r* implicit using the above formula.

The *bias* of an estimator 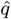 is defined as 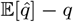. An unbiased estimator will on average return the correct value. In the case of mutation rate estimators, it is usually easier to derive the bias of 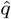 rather than 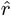 Even when there is a closed-form expression for the bias of 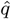, it does not lead to a closed-form expression for the bias of 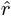, because 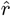 is not linear in 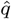. In this paper, we will derive the bias of our *q* estimators when possible. However, we will ultimately rely on experimental results to judge the bias of our estimators.

In this paper, we will take the *method-of-moments* approach to derive our estimators (Wasserman, 2013). We first decide on a random variable to observe, e.g. *I*^pp^ = |*sp*(*s*) ∩ *sp*(*t*)|. We then derive its expectation (possibly approximating it under some assumptions), e.g. 𝔼 [*I*^pp^] ≈ *L*(1 − *q*). We then take the observed value (denoted by 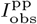 ), plug it into the expectation formula, and solve the formula for *q* to get the estimate. In our example, 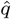 is the solution to the equation 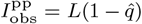, which is 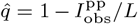.

## 3. The Presence-Presence Setting

In this section we consider the setting where the only thing we know about *s* and *t* are their spectra and *L*, i.e. no count information. We propose the following estimator

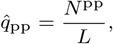

where *N*^pp^ = |*sp*(*t*) \ *sp*(*s*)| is the number of new distinct *k*-mers generated when *s* mutates into *t*. We will use the notation whereby a superscript of “pp” indicates the Presence-Presence setting, “pc” indicates the Presence-Count setting, and “cc” indicates the Count-Count setting. In this section, we explain how 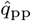 is derived and compare it to other known estimators for the Presence-Presence setting. We do not attempt to derive the bias of 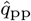, as it is technically complicated.

Consider the *k-span model*, which is to assume that

1. *s* does not contain any *k*-mer that occur more than once, and
2. new *k*-mers generated after mutations are distinct from each other and from the *k*-mers in *s*.

Let us define *I*^pp^ ≜ |*sp*(*s*) ∩ *sp*(*t*)| and use the shorthand *J* ≜ *J* (*sp*(*s*), *sp*(*t*)). In the *k*-span model, we have that 𝔼 [*I*^pp^] = *L*(1 − *q*) and the method-of-moments approach gives the estimator

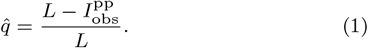

This estimator was introduced in Wu et al. (2025). Furthermore, in this model, *J* = *I*^pp^*/*(2*L* − *I*^pp^) and as a result we have that 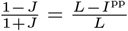. Thus, we can equivalently write

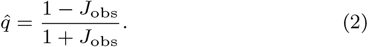

This is an improved version of the Mash estimator (Ondov et al., 2016), described in Sarmashghi et al. (2019) and in Appendix A.6 of Belbasi et al. (2022). Finally, in this model we have that 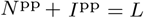, so we can equivalently write

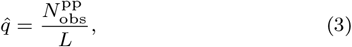

which corresponds the definition of our new estimator 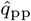. The three versions of 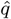 have the same derivation and are algebraically equivalent in the *k*-span model. However, when applied to data violating the *k*-span assumptions, we will see that three versions produce different results and 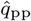 outperforms the others (Sec. 7).

The intuition for 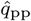 is based on considering what happens when assumption 1 is violated, i.e. there are repeats. Let *τ* be a *k*-mer with at least two occurrences in *s* and let *ν* be a *k*-mer that appears exactly once in *s*. Every mutation leads to a new *k*-mer in *N*^pp^, regardless of whether it happens in an occurrence of *τ* or *ν*. On the other hand, a mutation in one of the copies of *τ* does not effect *I*^pp^ while a mutation in *ν* increases *N*^pp^ by one. This makes Eq. 1 and Eq. 2, which rely on *I*^pp^, less accurate than Eq.3.

## 4. The Presence-Count Setting

In this section, we consider the Presence-Count setting and derive our estimator 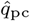 and its bias. Given two strings *s* and *t*, we define

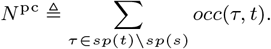

Intuitively, *N*^pc^ is the number of new *k*-mers generated when *s* mutates into *t*. We will derive 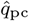 by applying a method-of-moments approach to *N*^pc^. Let us first define

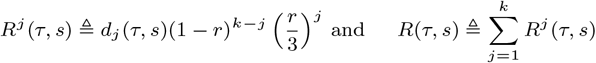

*R*^*j*^ (*τ, s*) is the probability that *τ* mutates to a *k*-mer that is in *s* and has a Hamming distance of *j* to *τ* . *R*(*τ, s*) is the probability that *τ* mutates to a *k*-mer that is in *s* but is different from *τ* . Note that we defined *R*^*j*^ and *R* in terms of *r* since it is visually clearer, but we can equivalently think of them as functions of *q*. We can now derive 𝔼 [*N*^pc^].

### Lemma 1

Let *s* be a string and let *t* be a string generated from *s* using the mutation process parameterized by *r*. Then

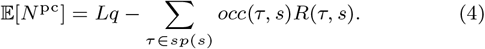

*Proof* Let *HD*(*τ, ν*) denote the Hamming distance between *k*-mers *τ* and *ν*. For 1 ≤ *i* ≤ *L*, let *s*_*i*_ be the *k*-mer starting at position *i* of *s*. Let *X*_*i*_ be an indicator random variable representing the event that *s*_*i*_ mutated to a *k*-mer that does not appear in *s*, i.e., *t*_*i*_ ∉ *s*. We can express *N*^pc^ as a sum *X*_*i*_s as follows.

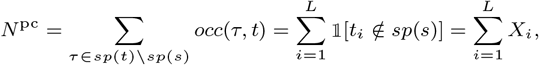

where 𝟙 is the indicator function for an event. Then,

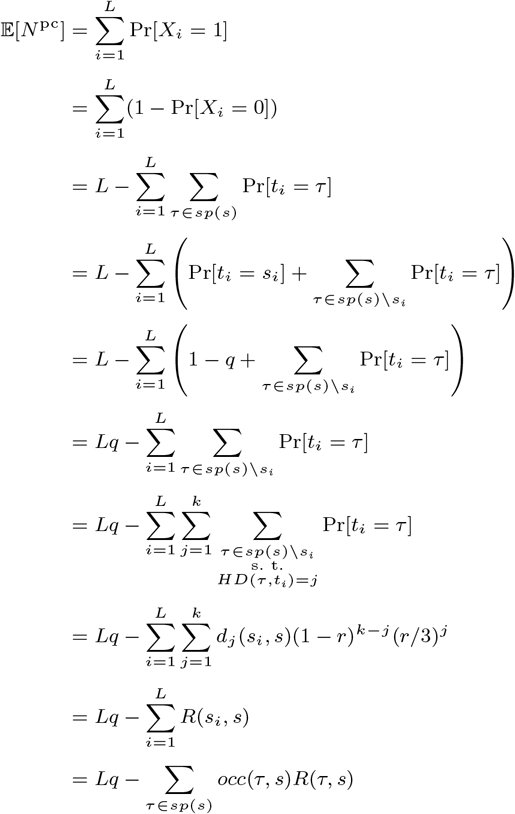

A straightforward method-of-moments estimator based on Eq. 4 would be to observe the value of *N*^pc^ (denoted as 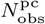 ), let 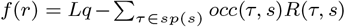, and use numerical methods to solve the equation 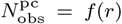 for *r*. However, there is no guarantee for the uniqueness of the solution. Furthermore, it would be time-consuming and superfluous to compute *R*(*τ, s*) for all *τ* .

Instead, we derive an estimator based on an approximation. Consider the two terms of Eq. 4. The first term is the expected number of positions whose *k*-mer mutated. This may over-count *N*^pc^, and the second term corrects this by accounting for the possibility that a position mutates but to something that is already in *s*. As we expect the first term to dominate, we approximate 𝔼 [*N*^pc^] ≈ *Lq*, leading to the estimator

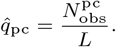

Unlike 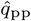, this formula accounts for the possibility that two occurrences of a *k*-mer in *s* mutate to the same new *k*-mer in *t*. The bias of 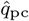 follows immediately from Lemma 1:

### Theorem 1

*The bias of* 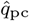 *is*

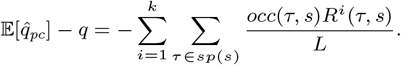

Note that the bias is always negative, meaning that, on average, 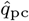 underestimates the true value. Moreover, the bias is a sum of *k* negative terms, which leads to the intuition of reducing the bias by including one of the terms inside the estimator itself—something we pursue in the next section.

## 5. The Count-Count Setting

In this section, we first present a novel Count-Count estimator 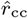 and then describe an alternative Count-Count estimator proposed by Rhie et al. (2020) in the context of assembly quality assessment.

### 5.1 New Estimator 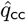

Let us build on 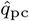 by trying to include the first term of the bias (i.e. − ∑_*τ*_ *occ*(*τ, s*)*R*^1^(*τ, s*)*/L*) into the estimator itself. With the method-of-moments approach, we would need to solve the following equation for *r*:

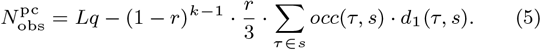

Unfortunately, we cannot solve this equation analytically and we cannot guarantee that a numerical method would produce a unique solution. Instead, we will first compute the 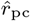 estimator and then plug into the right hand side of Eq.5. Our estimator is then defined as

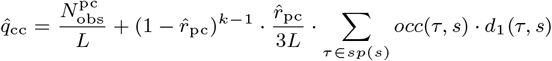

While this approach of plugging in one estimator in order to derive another one makes it challenging to prove anything, we will see that it performs extraordinarily well in practice.

To compute 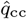, we need to compute *d*_1_(*τ, s*) for all *τ* ∈ *sp*(*s*). We do this in a straightforward two-pass algorithm using a hash table, resulting in runtime linear in *L*. There are more advanced ways of doing this (e.g. using suffix arrays), but we were not concerned with optimizing runtime since it was below one second.

### 5.2 Estimator mentioned in Rhie et al. (2020)

Rhie et al. (2020) mention an estimator that we recapitulate here to show how it fits in our framework. It is designed in the spirit of Eq. 1 by relying on the intersection size but also integrating count information. Consider the size of the weighted intersection between the *k*-mers of *s* and *t*:

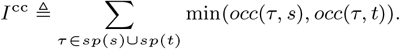

Let us assume that when a mutation occurs, the new *k*-mer is not in *s*. Before any mutations, *I*^cc^ = *L*. When a *k*-mer *τ* mutates to another *k*-mer *τ*^′^, it increases *occ*(*τ*^′^, *t*) by one and decreases *occ*(*τ, t*) by one. Since *τ*^′^ is not in *s, occ*(*τ*^′^, *t*) does not contribute anything to *I*^cc^. Therefore, the only effect of a mutation on *I*^cc^ is to decrease it by one. By linearity of expectation, we can add the probability of this happening for each of the *L* (non-distinct) *k*-mers and get that 𝔼 [*I*^cc^] ≈ *L*(1 − *q*). The method-of-moments approach then gives the estimator

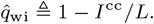

## 6. Combination With Sketching

A major reason why many estimators are based on *k*-mers rather than on the full sequences is that it makes them easily amenable to sketching. Sketching is a powerful technique that can make it possible to quickly compute all-pairs estimates on large datasets (Ondov et al., 2016). A *FracMinHash sketch sp*_*θ*_ (*s*) of a sequence *s* is defined as the subset of *sp*(*s*) that includes only the *k*-mers that map below a pre-defined threshold *θ*, under a fixed random hash function (Hera et al., 2023).

In this section, we will present a modification of 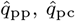 and 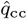 that work on data sketched with FracMinHash. Instead of observing *N*^pp^ or *N*^pc^, we now only observe them restricted to the sketched *k*-mers. Formally, let us define

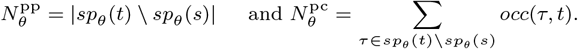

Intuitively, we expect each of these quantities to decrease by a factor of *θ* relative to their non-sketched versions. Based on this intuition, we define 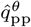 and 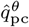 as 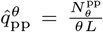 and 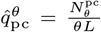. Formalizing our intuition, we prove that sketching does not effect the bias.

### Theorem 2

*The biases of* 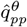 *and* 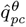

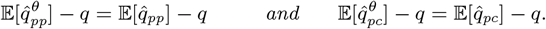

*Proof* First, we show that 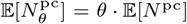. As before, let *X*_*i*_ be an indicator random variable representing the event that *s*_*i*_ mutated to a *k*-mer that does not appear in *s*. Let *Y*_*τ*_ be a binary random variable and it is 1 if *k*-mer *τ* hashes to less than *θ*. By the linearity, we have

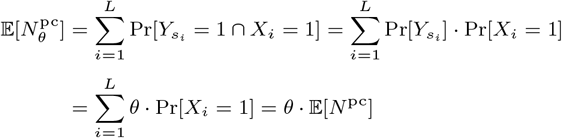

Then,

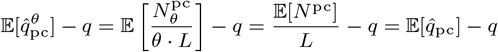

The proof for the bias of 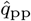 is similar and is omitted. □

Because 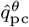 shows the same bias as 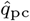, we use the same idea of Sec. 5.1 to obtain a stronger estimator 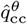 by partially correcting the bias of 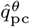, i.e.

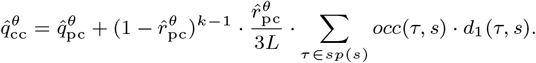

Note that ∑_*τ* ∈*s*_ *occ*(*τ, s*) · *d*_1_(*τ, s*)*/L* needs to be precomputed prior to sketching. It does not increase the space needed for the sketch as it becomes just a single constant that needs to be stored.

Because of the variance introduced by sketching, the observed quantity can exceed *θL* and then any of our three estimators can exceed 1. We therefore cap both estimators so that they never return greater than 1.

## 7. Empirical Results

We aim to evaluate the accuracy of our three novel estimators in relation to each other as well as to the other estimators mentioned in this paper. We also show an application to real data, where our estimators can be used to predict the ANI between various genomes. Our software is available on GitHub (Wu and Medvedev, 2025).

### 7.1. Datasets

We use four different base sequences with various levels of repetitiveness, summarized in Table S1. We use these sequences as representative examples spanning different levels of repetitiveness. Our main text evaluation is focused on the D-hardest sequence, which is a 100kbp-long sequences of alpha satellite DNA extracted from the human T2T chr21 centromere. We use a value of *k* = 30 for our evaluation on this sequence, as consistent with previous analyses (Wu et al., 2025), leading to 3, 987 distinct 30-mers. Over 70% of the distinct *k*-mers in D-hardest occur more than once and its *k*-mers have on average at least one other *k*-mer at a Hamming distance of one. D-hardest violates both assumption 1 (i.e. because it has repeats) and assumption 2 (i.e. because it has pairs of *k*-mers with small Hamming distances, a *k*-mer can mutate into one that is already in *s*). Since the differences between the estimators are more pronounced on this sequence, our main text focuses on D-hardest. The results on the three other sequences are presented in Sec. A of the Supplementary; they are all consistent with our findings on D-hardest but with less pronounced differences on less repetitive sequences.

### 7.2 Evaluation Metrics

We focus our evaluation on the estimators’ accuracy, measuring both their bias and variance. We do not perform a runtime or memory analysis because they each complete in less than a second in total on all of the four datasets and use negligible memory.

First, we benchmark each estimator on the four datasets using their default values of *k*, i.e. the values chosen in Wu et al. (2025) as most suitable for their analysis. We vary the mutation rate *r* from 0.001 to 0.251 and we show the distribution of 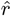 for 100 mutation-process simulation replicates for each *r* (e.g. Fig. 1). These experiments give a fine-grained separate view of the bias and variance.

**Fig. 1:**
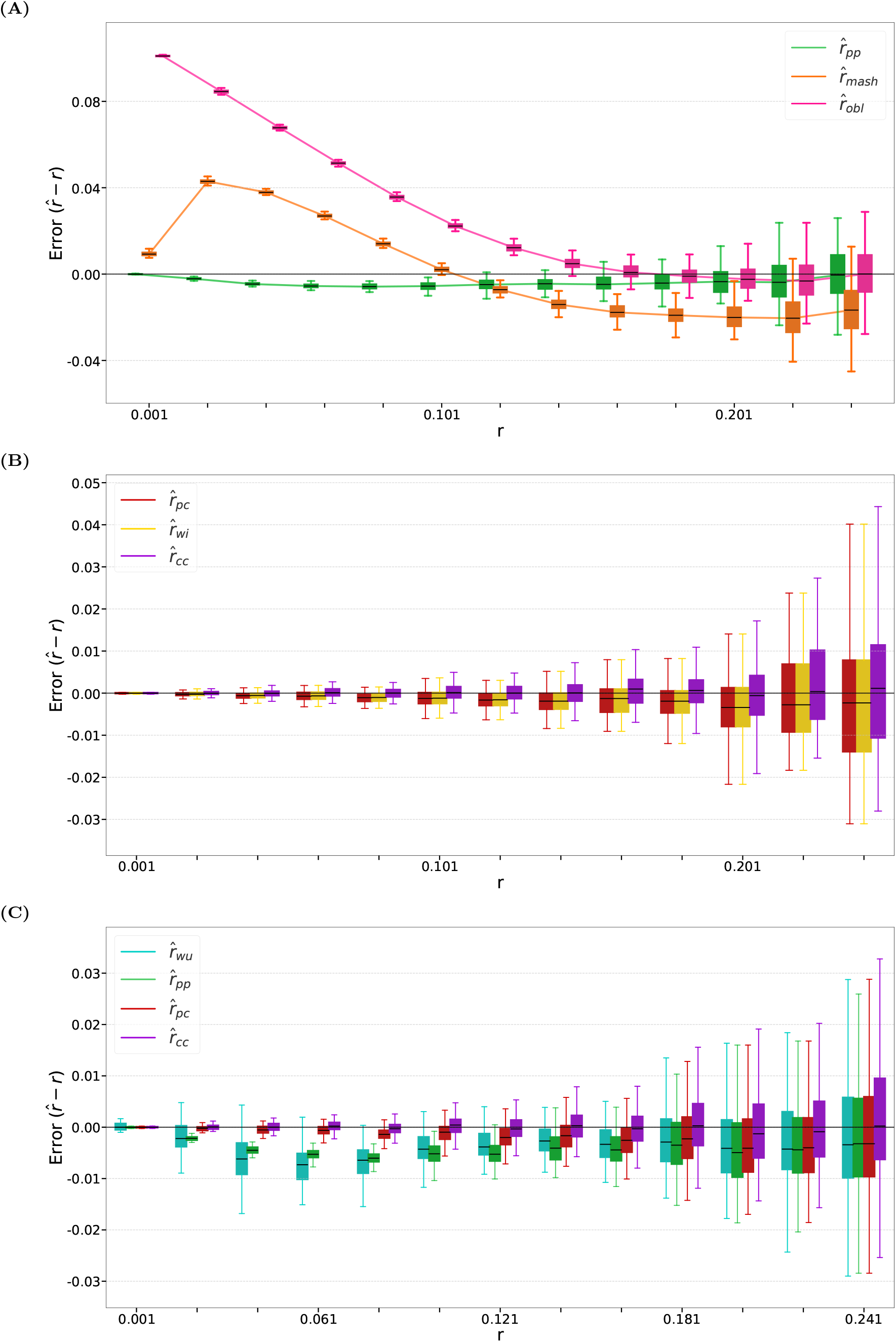
Comparison of estimator accuracy on D-hardest, with *k* = 30 and mutation rate *r* from 0.001 to 0.251 with step size 0.02. For each value of *r*, we show box plots over 100 mutation replicates. **(A)** Comparison of 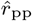 against other presence–presence estimators. **(B)** Comparison of 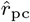 against 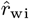and 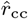. **(C)** Comparison of our three novel estimators against each other and against the 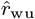 estimator of Wu et al. (2025).

Second, we vary both *k* and *r* and, for each (*k, r*) pair, compute the average relative absolute error: 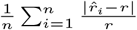(e.g. Fig. 2 and Table 2). We use *n* = 100 replicates for each (*k, r*) pair. This error combines the bias and variance into one metric, enabling us to easily visualize accuracy in two dimensions. Note that the two types of benchmarks emphasize different aspects of estimator performance.

**Table 2.** Errors of estimators under different parameter settings. Each cell shows the average relative absolute error of 100 replicates. Bold values highlight the lowest error in the respective class, while red/italic values highlight the lowest error overall. Underline denotes estimators we introduce in this paper. The 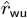 estimator falls in between the Presence-Presence and Presence-Count categories.

| Estimator | Category | $r = 0.001$ | | | $r = 0.010$ | | | $r = 0.100$ | | |
| --- | --- | --- | --- | --- | --- | --- | --- | --- | --- | --- |
| | | $k = 16$ | $k = 24$ | $k = 32$ | $k = 16$ | $k = 24$ | $k = 32$ | $k = 16$ | $k = 24$ | $k = 32$ |
| $\hat{r}_{obl}$ | Presence-Presence | 202 | 130 | 94 | 19 | 12 | 8.7 | 1.1 | .46 | .18 |
| $\hat{r}_{mash}$ | | 14 | 11 | 8.9 | 6.5 | 4.6 | 3.5 | .64 | .20 | **.013** |
| <u>$\hat{r}_{pp}$</u> |  | **.089** | **.083** | **.083** | **.14** | **.095** | **.073** | **.23** | **.097** | .044 |
| $\hat{r}_{wu}$ | | .88 | .73 | .67 | .30 | .20 | .20 | .37 | .13 | .037 |
| <u>$\hat{r}_{pc}$</u> | Presence-Count | .084 | .084 | .083 | .035 | .029 | .025 | .032 | .018 | .017 |
| $\hat{r}_{wi}$ | Count-Count | **.082** | **.084** | .081 | .030 | .027 | .024 | .028 | .017 | .017 |
| <u>$\hat{r}_{cc}$</u> |  | .083 | .085 | **.081** | **.027** | **.024** | **.023** | **.010** | **.011** | **.014** |

**Fig. 2:**
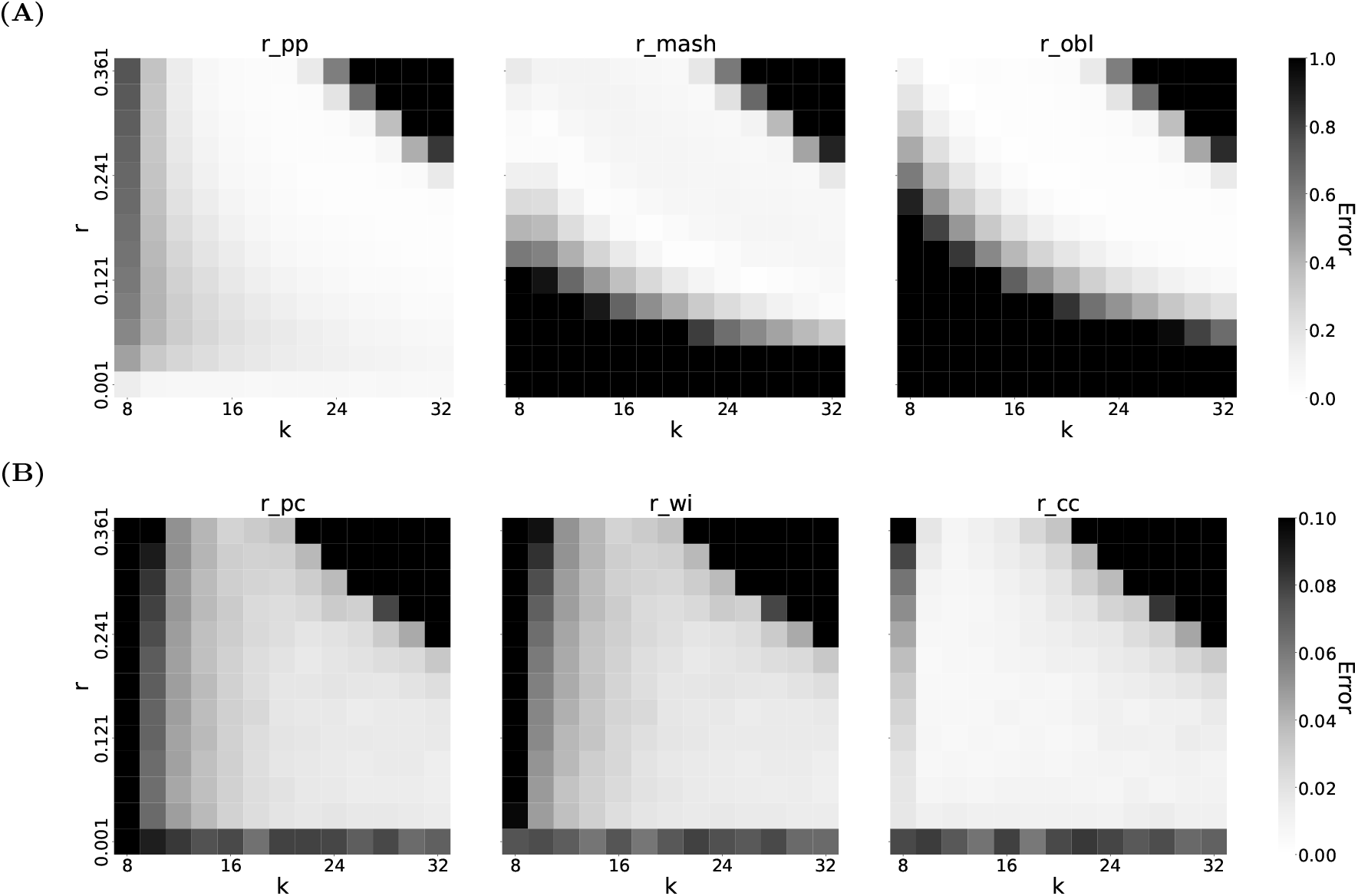
The estimators accuracy on D-hardest as a function of both *k* and *r*. Each cell shows the average relative absolute error of 100 replicates. Note that the scale of the heatmap is different between the top and bottom panels. Moreover, the errors are capped at 1.0 (for the panel A) and 0.10 (for the panel B), e.g., all errors greater than 1.0 are shown as 1.0 in the top panel.

As observed in our previous work (Wu et al., 2025), when *r* and/or *k* become sufficiently large, all *k*-mers mutate with very high probability, causing all tested estimators to return the value 1. We refer to this unstable behavior as *blow-up* and it manifests as high error in the top-right corner of the heatmaps.

### 7.3 Presence-Presence Setting

We compare 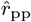 with the two other estimators in this category: the estimator defined by Eq. 1, which we refer to as 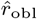, and the estimator defined by Eq. 2, which we refer to as 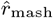 (this is the widely used Mash estimator with a binomial correction). Fig. 1A shows that for *k* = 30, 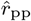 dominates 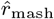across nearly all tested mutation rates; 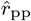 dominates 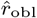 at lower values of *r* and has similar variance and bias for *r >* 0.161.

Fig. 2A presents heatmaps of estimator accuracy across a wide range of *k* and *r*. As expected, all estimators exhibit blow-up behavior for sufficiently large *k* and *r*. However, 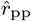 performs substantially better than 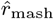 and 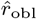 at smaller mutation rates.

### 7.4 Presence-Count and Count-Count Setting

For the Presence-Count setting, 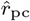 is the only estimator that we are aware of, while for the Count-Count setting, we have our new estimator 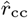 and the 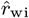 estimator. Fig. 1B shows the performance of these estimators with *k* = 30. and Fig. 2B shows the performance across a wide range of *k* and *r*. Table 2 shows specific values of errors under different settings of *k* and *r*.

First, we note that all estimators in these settings have smaller or similar bias and error than the 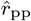 estimator, underscoring the general power of using count information.

Second, in the Count-Count setting, 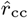 outperforms 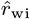for both *k* = 30 and more broadly across most tested (*k, r*) values. Although 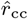 and 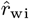 have access to the same *k*-mer count information, 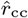 achieves nearly unbiased estimation for all tested values of *r* at *k* = 30.

Third, in the Presence-Count setting, 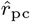 does not have a direct competitor, so we compare it against 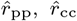, and 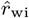. Compared to the 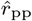 estimator of the Presence-Presence setting, 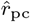 explicitly accounts for the event that multiple *k*-mers can mutate into the same novel *k*-mer, resulting in a smaller bias than 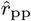. The estimator 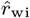 further addresses certain cases in which a *k*-mer mutates into another *k*-mer already present in the original sequence and, consequently, exhibits a slightly smaller bias than 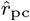for *r <* 0.051. Nevertheless, 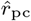and 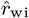 have very similar performance. Compared to 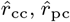 does not perform as well, which is expected since 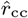 is designed to specifically offset some of the bias of 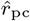.

### 7.5 Comparison Against Estimator From Wu et al. (2025)

In a previous work, we tackled a similar problem and developed a single repeat-robust estimator (Wu et al., 2025). We will refer to it as 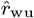 here. It was the first *k*-mer-based estimator evaluated for accuracy in highly repetitive settings, showing robustness in these settings compared to 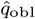. It does not neatly fit into the categories here, as it uses the abundance histogram of *s*, i.e. the histogram of *k*-mer occurrence counts. While this is based on the count of the *k*-mers of *s*, it is only a summary and can be approximated using a related species or a related type of sequence. In our framework, therefore, 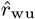 lies somewhere between the Presence-Presence and the Count-Presence setting. Fig. 1C compares 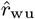 against our three estimators for *k* = 30. Table 2 and Fig. S1 show the error across different settings of *k* and *r*.

Despite relying on less information, 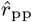is overall a better estimator than 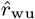. The relative bias depends on 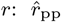has slightly smaller bias for 0.001 ≤ *r* ≤ 0.091 and has slightly larger bias for 0.101 ≤ *r* ≤ 0.231 (Fig. 1C). However, 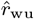 often has more variance than 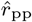, as reflected in both Fig. 1C and the bigger relative absolute errors shown in Table 2 and Fig. S1.

The improvement of 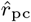 over 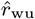 is more stark, as 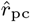 consistently has lower bias than 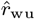 (Fig. 1C) and has consistently lower overall error across all values of *k* and *r* (Table 2 and Fig. S1). The superior performance of 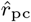 relative to 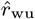 highlights the benefits of accounting for novel *k*-mers in the estimate formula, as 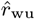 does not take into account any *k*-mers in *t* that are not in *s*.

### 7.6 Combination With Sketching Techniques

In Sec. 6, we proved that sketching does change the bias of 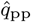 and 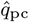. Here, we evaluate empirically the bias of all our three estimators. Fig. S2 shows the performance of 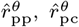, and 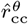 using *θ* ∈ {0.1, 0.01}. We observe that sketching does not introduce any systematic bias, in any of the three estimators. As expected with sketching, we observe increased variance for *θ* = 0.1 and even larger variance for *θ* = 0.01. Overall, our results confirm that our estimators can be applied naturally to sketched data, with the obvious caveat that smaller sketches will lead to larger variance of the estimator.

### 7.7 ANI Estimation on Real Genomes

To evaluate the applicability of our estimators on real data, we use 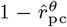 and 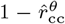to estimate ANI between real genomes. We use the slow but accurate alignment-based OrthoANIu (Yoon et al., 2017) to compute the ground truth, consistent with previous work (Shaw and Yu, 2023).

We evaluate using a dataset from Hera et al. (2023). They first chose ten representative genomes from the Genome Taxonomy Database, including seven bacterial and three archaeal species. For each representative genome, they extract additional genomes from along the evolutionary path to the root, selecting an additional three non-representative genomes at each taxonomic rank on the path up the tree. They construct pairs for comparison by matching each representative genome with the genomes selected along its evolutionary path up the tree. We further filtered out three pairs that had accession numbers that were not currently available. The resulting dataset contains 189 pairs and covers a wide range of ANI from ∼ 60% to 100%.

We benchmark our estimators against those of of Mash (Ondov et al., 2016), sourmash (Irber et al., 2024), FastANI (Jain et al., 2018), and skani (Shaw and Yu, 2023). FastANI and skani rely on mapping techniques with seeds; we run them with default parameters. Sourmash and Mash are both similar to our estimators in that they rely on *k*-mer sketches. We therefore set *k* = 19 for our tools and sourmash and Mash. Sourmash also uses FracMinHash, so we set *θ* = 0.01 for both sourmash and our estimators. Mash uses MinHash sketching instead of FracMinHash, so we set the size of its sketch to be 30,000 to roughly match the sketch sizes obtained with FracMinHash of *θ* = 0.01. Aside from this, we use the default parameters for sourmash and Mash. Table S2 in the Supplementary gives the full commands and data used.

Fig. S3 shows the estimator results relative to the ground truth. For some pairs at ANI *<* 85%, some of the estimators either do not report an estimate or report 0; we refer to these as *uncomputable* pairs. In order to make a fair comparison, we measure both the number of uncomputable pairs and the accuracy of the predictions at ANI *>* 85% (Table 3).

**Table 3.**
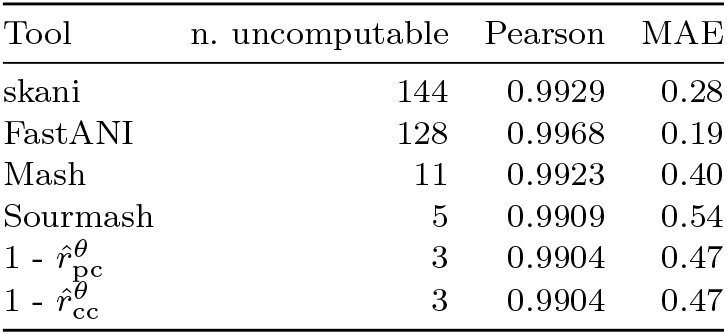
Results on the ANI benchmark. The number of uncomputable pairs is the number of pairs for which the estimator either returns 0 or does not return anything. Pearson R value (Pearson) and mean absolute error (MAE) are shown for a subset of pairs with ANI*>* 85%, to avoid penalizing for uncomputable pairs.

| Tool | n. uncomputable | Pearson | MAE |
| --- | --- | --- | --- |
| skani | 144 | 0.9929 | 0.28 |
| FastANI | 128 | 0.9968 | 0.19 |
| Mash | 11 | 0.9923 | 0.40 |
| Sourmash | 5 | 0.9909 | 0.54 |
| $1 - \hat{r}_{pc}^\theta$ | 3 | 0.9904 | 0.47 |
| $1 - \hat{r}_{cc}^\theta$ | 3 | 0.9904 | 0.47 |

The most accurate estimators at high ANI levels are skani and FastANI; however, they are not able to estimate ANI for more than 100 pairs at lower ANI levels. On the other hand, our estimators are the most comprehensive and are able to compute all except three pairs. Overall, there is a general pattern that more comprehensive estimators are less accurate at high ANI levels, making the choice of the best estimator a trade-off.

## 8. Discussion and Conclusion

In this paper, we studied the problem of estimating the mutation rate of a process that transforms an arbitrary string *s* into a string *t* by introducing substitutions at rate *r*. We focused on estimators that are based on *k*-mers, as they can easily be combined with sketching to make them scalable to large datasets. We observed that various estimators for this problem, including our own, can be categorized according to whether they have access to *k*-mer counts or to only presence/absence information. We presented three novel estimators, 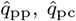, and 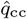, summarized in Table 1. On highly repetitive data, 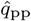 and 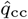 perform best in their category, with 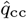 outperforming all tested estimators in all categories. The 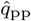 estimator is desirable when no count information is available, while the 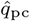 estimator offers a middle ground trade-off between accuracy and the amount of information required from the original and mutated sequences.

The main insight behind our 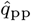 and 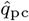 estimators is that in a repeat setting, it is important to count the *k*-mers that are newly created in *t* and do not appear in *s*. We reflect this in the title, i.e., *novel k*-mers are a *gift* that we must make use of. In many cases, such as 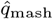 and 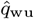, the formula relies on the observed number of shared *k*-mers. This works fine in the absence of repeats or mutations resulting in spurious matches, as there is a one-to-one correspondence between *k*-mers removed from the intersection and newly created *k*-mers. However, when there are repeats, a mutation in a repetitive *k*-mer *τ* only destroys one copy of *τ* and does not remove *τ* from the count of shared *k*-mers. On the other hand, it does add to the count of new *k*-mers, as *τ*^′^ is added as a new *k*-mer to *t*. We show that these types of events hurt the performance 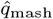 and 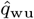, though the effect on 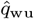is compensated by its use of counts. Our 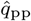 and 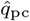 estimators, on the other hand, are fully based on the count of new *k*-mers, giving them a performance advantage.

In the Count-Count setting, 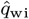 already accounts for this insight; in order to out-perform it, we add some accounting for the possibility that a *k*-mer mutates into a *k*-mer that is already in *s*. Consider the possibility that a *k*-mer *τ* in *s* mutates to a *k*-mer *ν* while a *k*-mer *ν* in *s* mutates into *τ* . The likelihood of this is not directly related to repeats, as it can happen in a repeat-free genome; it is related to two *k*-mers in *s* having a small Hamming distance between them. The 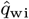 estimator does not model the chance of this event happening. Our estimator 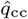 models this event happening for the case that the Hamming distance is one; we believe this accounts for its improved performance.

Our mutation model is naturally an idealized version of reality and does not explicitly account for factors such as ploidy or indels. Nevertheless, our estimators can be used as part of downstream tools. For example, the Merqury tool (Rhie et al., 2020) computes the quality of a candidate assembly by comparing its *k*-mer content to the *k*-mer content of the sequencing data used to validate it. The score that Merqury reports is closely related to our 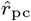estimator, as it essentially treats the assembly as a mutated string *t* from the original genome *s*. The *k*-mer counts of *t* are given by the assembly; to approximate the presence-absence spectrum of *s*, Merqury uses higher-copy *k*-mers from the read set. Note that what we are describing here differs from the 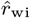estimator, which is also alluded to in Rhie et al. (2020).

A natural question arising from our work is how to choose *k* and *θ*. Guidance on selecting *k* is partially addressed by our heatmaps, which display estimator accuracy across a grid of (*k, r*) values for both repetitive and typical genomic settings. These plots allow users to identify the stable operating regime for their sequence type. Additionally, in our previous work (Wu et al., 2025), we introduced *P*_empty_ as a heuristic stability criterion: given sequence length *L*, mutation rate *r*, and *k, P*_empty_ quantifies the probability that a *k*-mer has no surviving copies after mutation, providing a principled threshold below which the estimator is expected to be unstable. The choice of *θ* (the sampling rate for sketching) has been widely studied in the sketching literature, though not for our new estimators in particular. Our analysis here (e.g. Fig. S2) can give a rough guidance for choosing *θ*, but a more rigorous analysis of sketch estimator stability in repetitive regions remains an interesting problem.

Our insights open the door for the continued improvement of estimators. One particular direction is the Count-Presence setting which we have not discussed in this paper. Note that this setting is not symmetric to the Presence-Count setting; e.g. the counts in *s* are a deterministic variable while the counts in *t* are a random variable. It remains an open problem to determine if estimators in the Count-Presence setting can outperform estimators in the Presence-Count setting. As more repetitive sequences become available, we expect future work to uncover new insights to further drive the improved quality of mutation rate estimators.

## Acknowledgments

We thank Antonio Blanca for useful discussions. This material is based upon work supported by the National Science Foundation under Grants No. DBI2138585 and OAC1931531. Research reported in this publication was supported by the National Institute Of General Medical Sciences of the National Institutes of Health under Award Number R01GM146462. The content is solely the responsibility of the authors and does not necessarily represent the official views of the National Institutes of Health.

## A. Supplementary information for “The gift of novelty: repeat-robust *k*-mer-based estimators of mutation rates” by Haonan Wu and Paul Medvedev

**Table S1.**
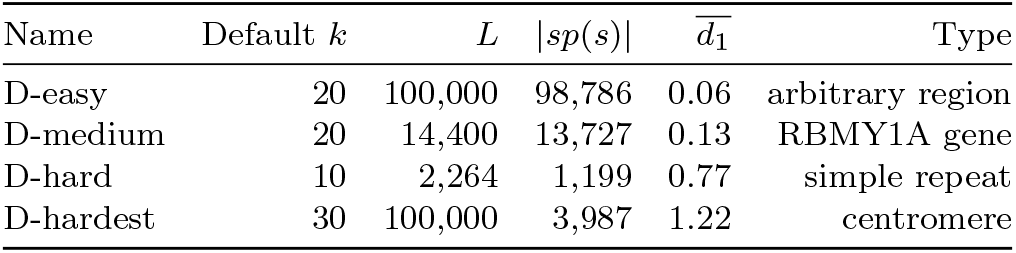
Sequence properties of our four datasets. These sequences were originally used for evaluation by Wu et al. (2025). They are extracted from the human T2T-CHM13v2.0 reference and made available to download directly https://zenodo.org/records/18303511. The 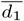 column shows the average number of neighbors that have a Hamming distance of 1, i.e. 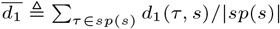.

| Name | Default $k$ | $L$ | $\vert sp(s)\vert $ | $\bar{d}_1$ | Type |
| --- | --- | --- | --- | --- | --- |
| D-easy | 20 | 100,000 | 98,786 | 0.06 | arbitrary region |
| D-medium | 20 | 14,400 | 13,727 | 0.13 | RBMY1A gene |
| D-hard | 10 | 2,264 | 1,199 | 0.77 | simple repeat |
| D-hardest | 30 | 100,000 | 3,987 | 1.22 | centromere |

**Table S2.**
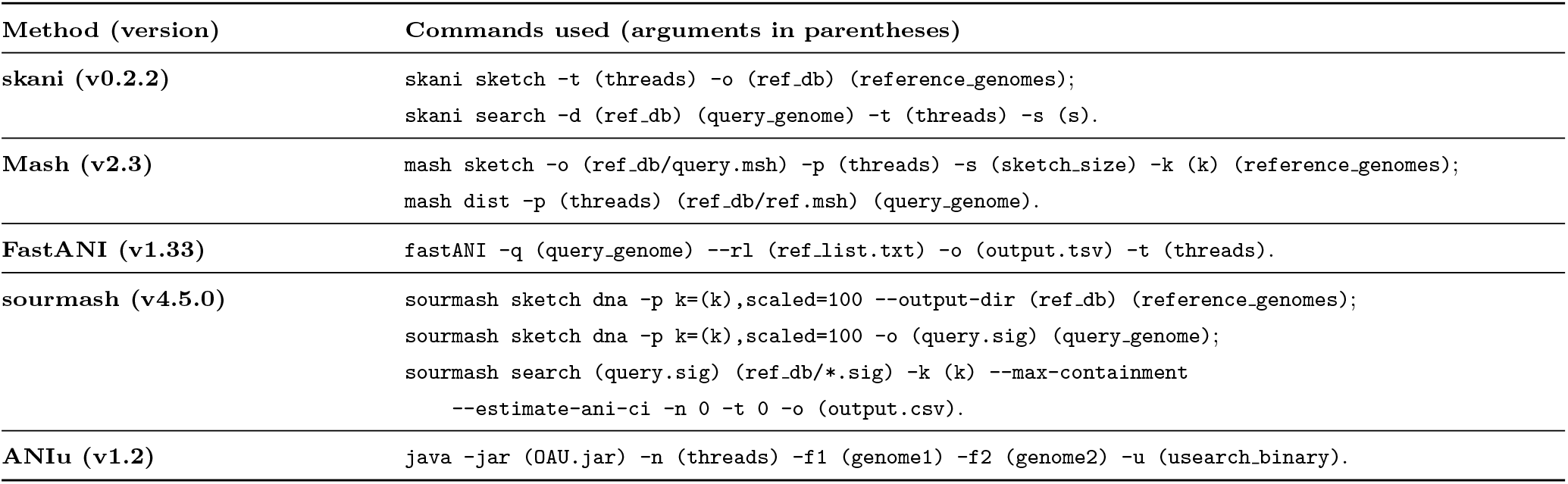
Commands used in our ANI benchmark pipeline. The dataset is available for download at https://github.com/bluegenes/2022-focused-cani-comparisons/blob/main/gtdb-rs207.common-sp10-evolpaths.csv.

**Fig. S1:**
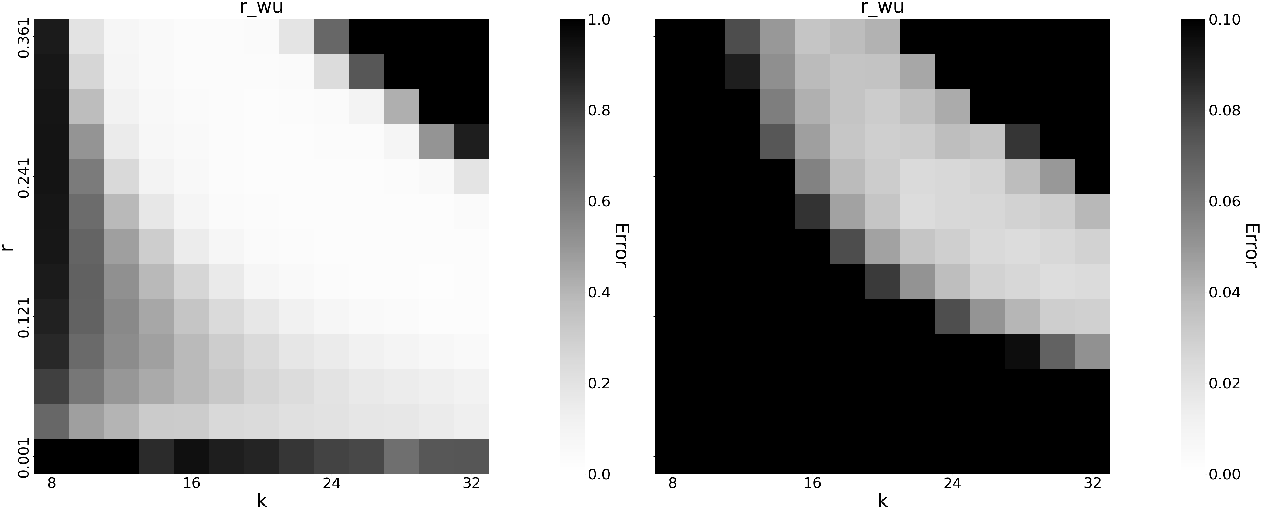
Heatmaps showing the performance of the 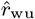 estimator on D-hardest. The left heatmap shows the error using a scale up to 1.0, and the right heatmap shows the error using a finer resolution of a scale up to 0.1.

**Fig. S2:**
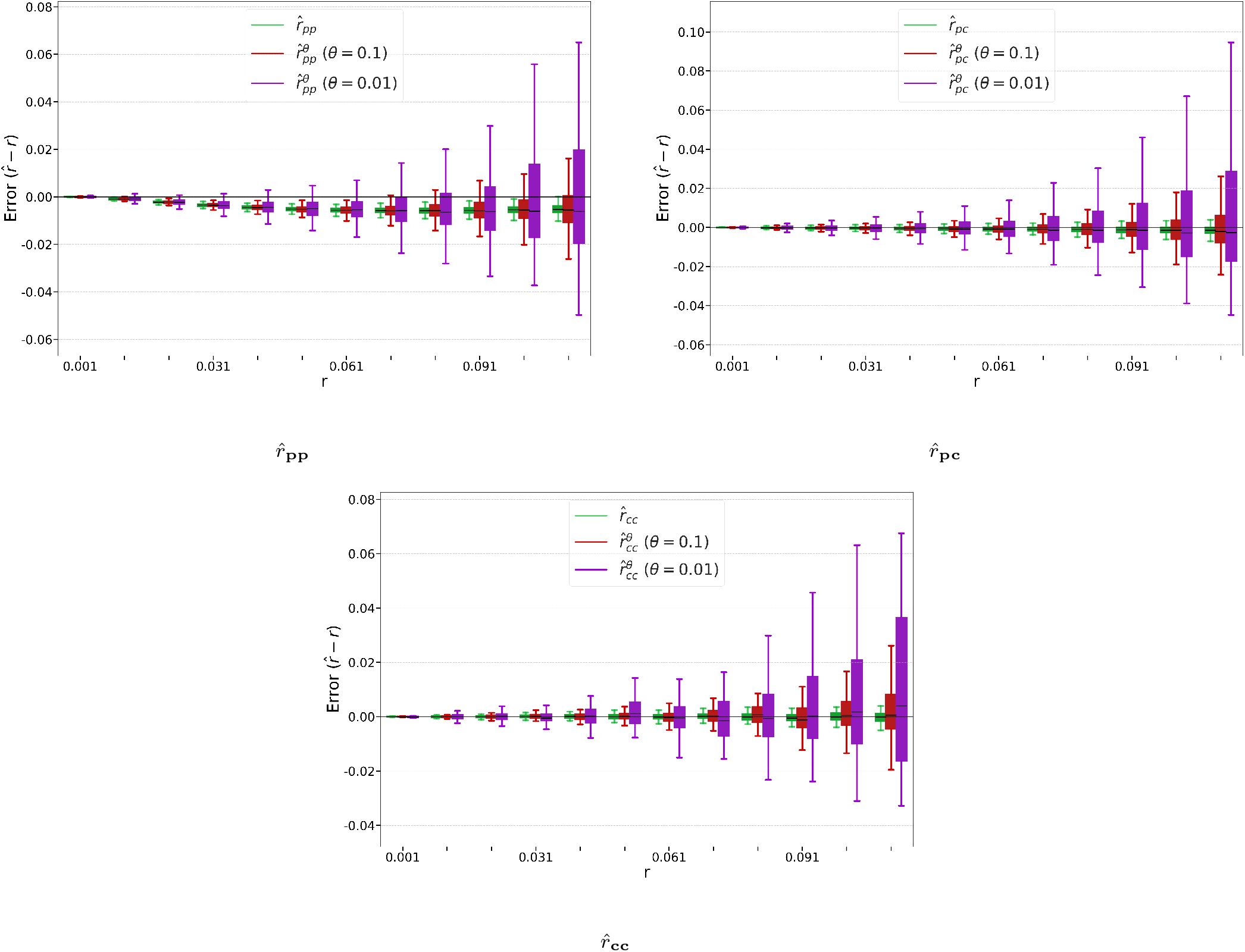
Effect of sketching on the accuracy of our three estimators, on D-hardest. Top left is 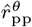, top right is 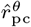, and bottom is 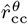 . We vary mutation rates in *r* ∈ [0.001, 0.111], using a step size of 0.01. Because sketching introduces additional variance, estimator blow-up occurs at smaller values of *r*, and we therefore restrict our evaluation to *r* ≤ 0.111.

**Fig. S3:**
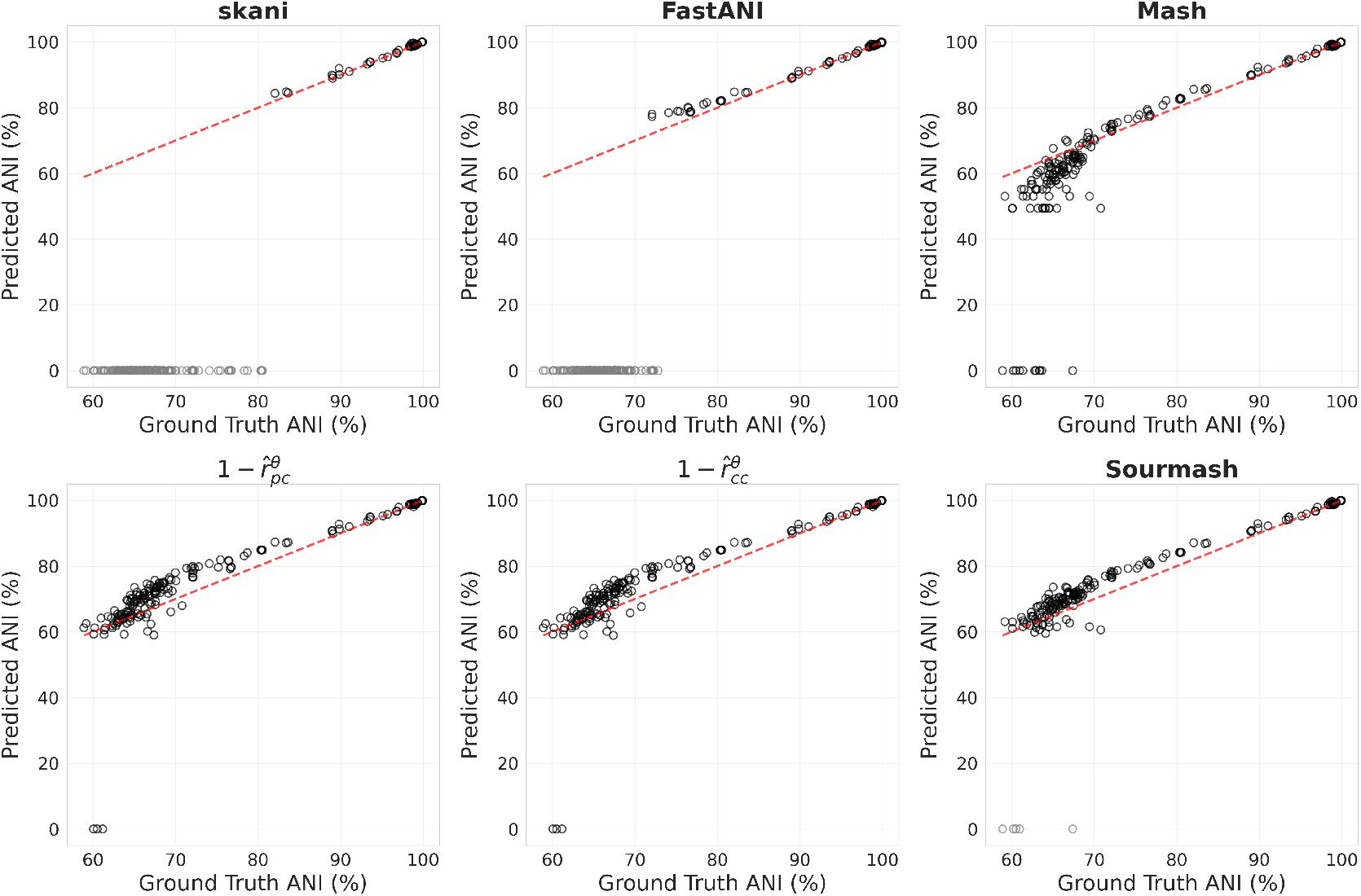
Performance of various estimators on the ANI benchmark. The dotted line represents a perfect predictor. For genome pairs where the estimator was unable to compute a value, we show the predicted value as 0.

### A.1. Comparison of all the estimators on D-easy, D-med, and D-hard

In the main text, we focused on the evaluation of the D-hardest dataset. For completeness, we also include this Supplementary section to show the results on the three easier datasets: D-easy, D-med, D-hard. These datasets were first introduced by Wu et al. (2025) to have progressive levels of repetitiveness. D-easy is an arbitrarily chosen substring of chr6, with less than 1% of *k*-mers being non-singleton. D-med is the sequence of the chrY RBMY1A1 gene, with approximately 3% of *k*-mers being non-singleton. D-hard is a subsequence of D-med that is annotated as a simple repeat, with more than 40% of *k*-mers being non-singletons. These datasets are summarized in Table S1. Fig. S4 shows the results for a single *k* value and The results are consistent with our findings for D-hardest but the differences become less pronounced on less repetitive sequences. For D-easy and D-med, all the estimators perform relatively well though 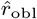 and 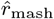show more bias than the rest. For D-hard, the relative performance is similar to that on D-hardest. Fig. S5 shows the heatmaps for a wide range of *k* and *r*.

**Fig. S4:**
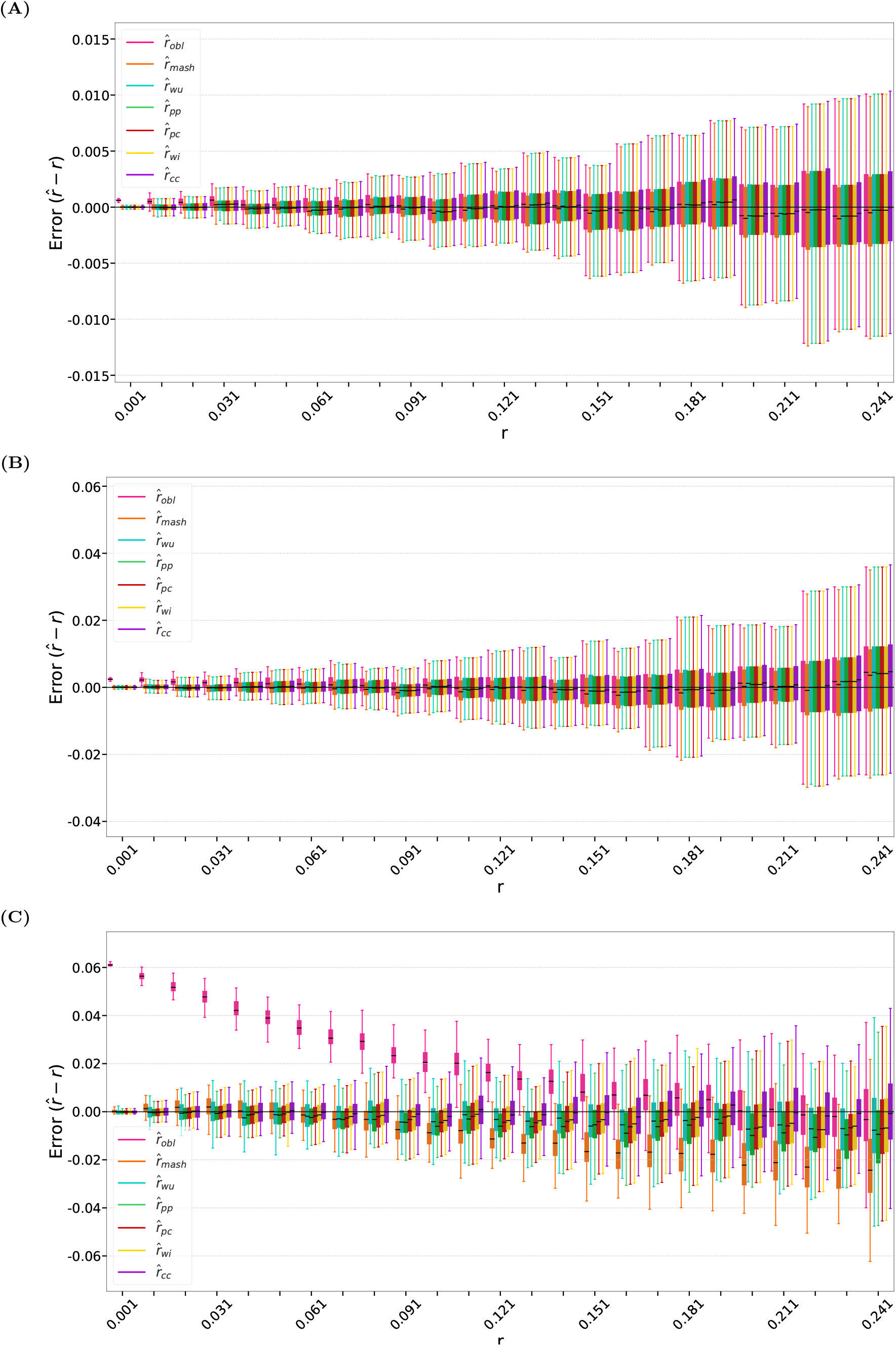
Comparison of all estimators on the **(A)** D-easy, **(B)** D-med, and **(C)** D-hard datasets. The *k* values used are as shown in Table S1, i.e. for D-easy, *k* = 20, for D-med, *k* = 20, and for D-hard, *k* = 10.

**Fig. S5:**
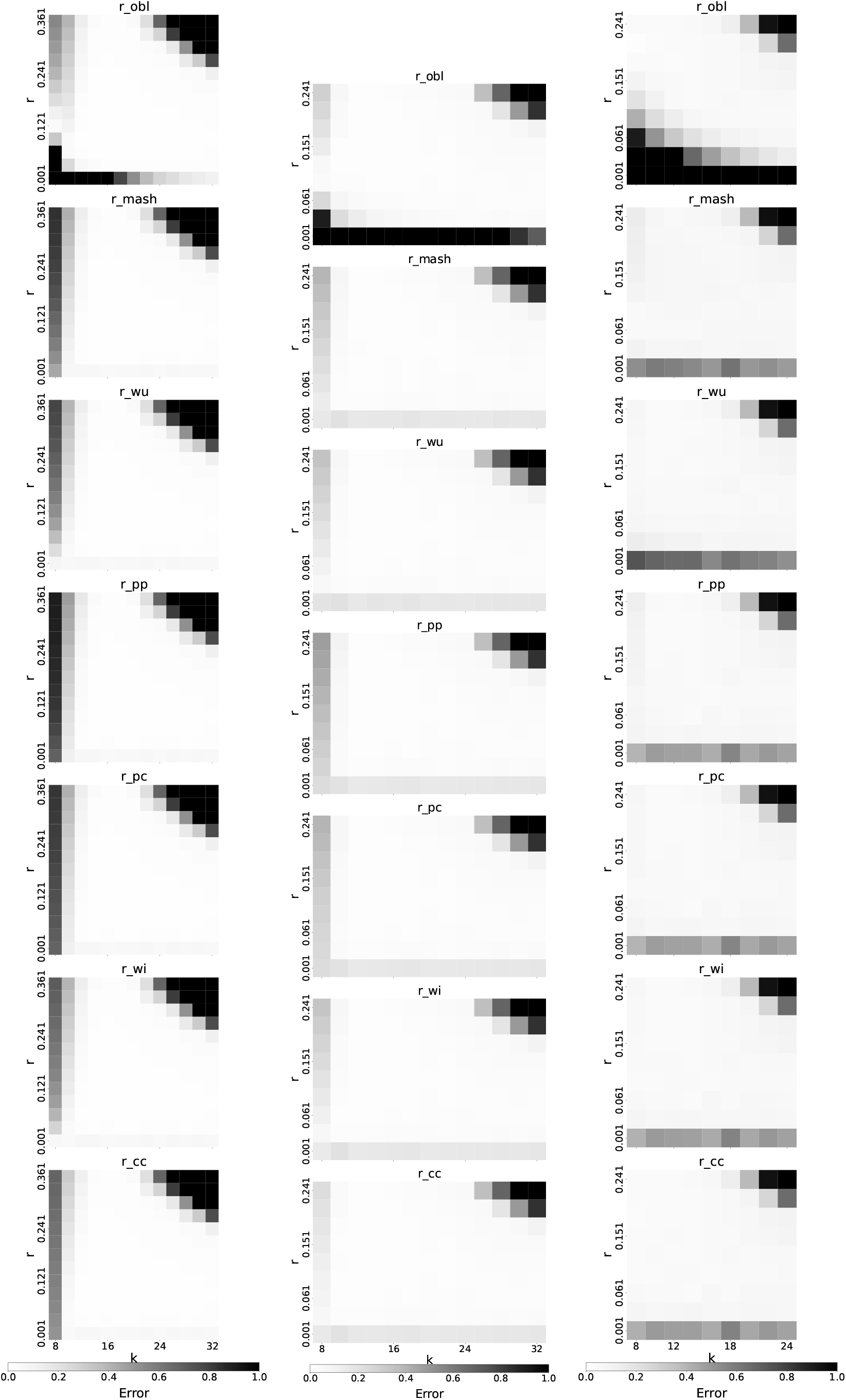
Heatmap comparison of all estimators on the D-easy (left column), D-med (middle column), and D-hard datasets (right column).

## References

Belbasi, M., Blanca, A., Harris, R. S., Koslicki, D., and Medvedev, P. (2022). The minimizer Jaccard estimator is biased and inconsistent. Bioinformatics, 38(Supplement 1):i169–i176.

Blanca, A., Harris, R. S., Koslicki, D., and Medvedev, P. (2022). The statistics of k-mers from a sequence undergoing a simple mutation process without spurious matches. Journal of Computational Biology, 29(2):155–168.

Dayhoff, M. O. (1969). Computer analysis of protein evolution. Scientific American, 221(1):86–95.

Durbin, R., editor (2013). Biological sequence analysis. Cambridge Univ. Press, Cambridge [u.a.], 17. print. edition.

Henikoff, S. and Henikoff, J. G. (1992). Amino acid substitution matrices from protein blocks. Proceedings of the National Academy of Sciences, 89(22):10915–10919.

Hera, M. R. and Koslicki, D. (2025). Estimating similarity and distance using FracMinHash. Algorithms for Molecular Biology, 20(1):1–13.

Hera, M. R., Liu, S., Wei, W., Rodriguez, J. S., Ma, C., and Koslicki, D. (2024). Metagenomic functional profiling: to sketch or not to sketch? Bioinformatics, 40(Supplement 2):ii165–ii173.

Hera, M. R., Medvedev, P., Koslicki, D., and Blanca, A. (2025). Estimation of substitution and indel rates via k-mer statistics. In 25th International Conference on Algorithms for Bioinformatics (WABI 2025), volume 344, pages 16:1–16:15. Schloss Dagstuhl – Leibniz-Zentrum für Informatik.

Hera, M. R., Pierce-Ward, N. T., and Koslicki, D. (2023). Deriving confidence intervals for mutation rates across a wide range of evolutionary distances using FracMinHash. Genome research, 33(7):1061–1068.

Irber, L., Pierce-Ward, N. T., Abuelanin, M., Alexander, H., Anant, A., Barve, K., Baumler, C., Botvinnik, O., Brooks, P., Dsouza, D., et al. (2024). sourmash v4: A multitool to quickly search, compare, and analyze genomic and metagenomic data sets. Journal of Open Source Software, 9(98):6830.

Jain, C., Rodriguez-R, L. M., Phillippy, A. M., Konstantinidis, K. T., and Aluru, S. (2018). High throughput ani analysis of 90k prokaryotic genomes reveals clear species boundaries. Nature communications, 9(1):1–8.

Logsdon, G. A., Rozanski, A. N., Ryabov, F., Potapova, T., Shepelev, V. A., Catacchio, C. R., Porubsky, D., Mao, Y., Yoo, D., Rautiainen, M., et al. (2024). The variation and evolution of complete human centromeres. Nature, 629(8010):136–145.

Ondov, B. D., Treangen, T. J., Melsted, P., Mallonee, A. B., Bergman, N. H., Koren, S., and Phillippy, A. M. (2016). Mash: fast genome and metagenome distance estimation using MinHash. Genome Biology, 17(1):132.

Rathore, S. P. S. and Kashyap, N. (2026). Estimators for substitution rates in genomes from read data. arXiv preprint arXiv:2601.07546.

Rhie, A., Walenz, B. P., Koren, S., and Phillippy, A. M. (2020). Merqury: reference-free quality, completeness, and phasing assessment for genome assemblies. Genome Biology, 21(1).

Sarmashghi, S., Bohmann, K., Gilbert, M. T. P., Bafna, V., and Mirarab, S. (2019). Skmer: assembly-free and alignment-free sample identification using genome skims. Genome Biology, 20(1):1–20.

Shaw, J. and Yu, Y. W. (2023). Fast and robust metagenomic sequence comparison through sparse chaining with skani. Nature Methods, 20(11):1661–1665.

Shaw, J. and Yu, Y. W. (2024). Rapid species-level metagenome profiling and containment estimation with sylph. Nature Biotechnology, pages 1–12.

Song, K., Ren, J., Reinert, G., Deng, M., Waterman, M. S., and Sun, F. (2014). New developments of alignment-free sequence comparison: measures, statistics and next-generation sequencing. Briefings in bioinformatics, 15(3):343–353.

Wasserman, L. (2013). All of statistics: a concise course in statistical inference. Springer Science & Business Media.

Wu, H., Blanca, A., and Medvedev, P. (2025). A *k*-mer-based estimator of the substitution rate between repetitive sequences. In 25th International Conference on Algorithms for Bioinformatics (WABI 2025), volume 344 of Leibniz International Proceedings in Informatics (LIPIcs), pages 20:1–20:20.

Wu, H. and Medvedev, P. (2025). Accurate repeat-aware kmer based estimator. https://github.com/medvedevgroup/Accurate_repeat-aware_kmer_based_estimator.

Yoon, S.-H., Ha, S.-m., Lim, J., Kwon, S., and Chun, J. (2017). A large-scale evaluation of algorithms to calculate average nucleotide identity. Antonie Van Leeuwenhoek, 110(10):1281– 1286.

Zielezinski, A., Vinga, S., Almeida, J., and Karlowski, W. M. (2017). Alignment-free sequence comparison: benefits, applications, and tools. Genome biology, 18(1):186.

